# Delineating yeast cleavage and polyadenylation signals using deep learning

**DOI:** 10.1101/2023.10.10.561764

**Authors:** Emily Kunce Stroup, Zhe Ji

## Abstract

3’-end cleavage and polyadenylation is an essential process for eukaryotic mRNA maturation. In yeast species, the polyadenylation signals that recruit the processing machinery are degenerate and remain poorly characterized compared to well-defined regulatory elements in mammals. Especially, recent deep sequencing experiments showed extensive cleavage heterogeneity for some mRNAs in *Saccharomyces cerevisiae* and uncovered the polyA motif differences between *S. cerevisiae* vs. *Schizosaccharomyces pombe*. The findings raised the fundamental question of how polyadenylation signals are formed in yeast. Here we addressed this question by developing deep learning models to deconvolute degenerate *cis*-regulatory elements and quantify their positional importance in mediating yeast polyA site formation, cleavage heterogeneity, and strength. In *S. cerevisiae*, cleavage heterogeneity is promoted by the depletion of U-rich elements around polyA sites as well as multiple occurrences of upstream UA-rich elements. Sites with high cleavage heterogeneity show overall lower strength. The site strength and tandem site distances modulate alternative polyadenylation (APA) under the diauxic stress. Finally, we developed a deep learning model to reveal the distinct motif configuration of *S. pombe* polyA sites which show more precise cleavage than *S. cerevisiae*. Altogether, our deep learning models provide unprecedented insights into polyA site formation across yeast species.

## INTRODUCTION

In eukaryotes, the 3’-ends of mRNAs are co-transcriptionally generated by a two-step catalytic process: endonucleolytic cleavage followed by polyA tail synthesis (Richard and Manley 2009). Cleavage and polyadenylation happens through the recruitment of processing factors to sequence motifs around polyA sites. While polyA signals in mammals are well-defined, the sequence composition of yeast polyA sites remains poorly understood despite some processing factors demonstrating homology across the species (Mandel et al. 2008; Tian and Graber 2012). In mammals, the essential motifs include the CPSF binding site (AAUAAA or close variants) typically located ∼20 nt upstream of the cleavage sites and the CSTF binding site (U/GU-rich motifs) ∼20 nt downstream. Additional motifs can play auxiliary regulatory roles. However, the polyA signals in *Saccharomyces cerevisiae* are highly degenerate and variable across genes.

Previous experimental and computational characterization of yeast polyA sites identified *cis*-elements promoting site formation: the UA-rich elements located 40 nt upstream (designated as the efficiency element) bound by the cleavage and polyadenylation factor 1B (CF1B), the A-rich motif 20 nt upstream (named as the positioning element) bound by CF1A, and U-rich elements surrounding the cleavage site bound by the polyadenylation factor complex CPF (Guo and Sherman 1996; Graber et al. 1999a; Graber et al. 1999b). The composition of the polyA site sequence itself can also influence cleavage site selection (Guo and Sherman 1996; Graber et al. 1999a; Graber et al. 1999b; Dichtl and Keller 2001). However, some binding motifs (e.g., UA-rich and A-rich elements) can be highly degenerate (Guo and Sherman 1996). Currently, there are no quantitative comparisons of the relative importance of near-cognate polyA signals.

Recent advances in mapping 3’-ends of RNA isoforms in *S. cerevisiae* using deep sequencing showed that the cleavage heterogeneity is quite extensive for some mRNAs (Moqtaderi et al. 2013). A cleavage zone can span dozens of nucleotides in a yeast mRNA, which rarely happens for mammalian genes. It remains unknown if the polyadenylation events occurring across a cleavage zone represent the site micro-heterogeneity or the utilization of separate alternative polyA sites. Both heterogeneous cleavage and alternative polyadenylation (APA) can generate mRNAs with different 3’UTR regulating subcellular RNA localization, translation efficiency, and stability (Tian and Manley 2017; Gruber and Zavolan 2019; Mitschka and Mayr 2022). mRNA isoforms with only a few nucleotide differences in 3’-ends can show drastic variability in RNA secondary structure and stability (Geisberg et al. 2014; Moqtaderi et al. 2018). Extensive APA regulation occurs during the stress response (Graber et al. 2013; Geisberg et al. 2020). Studies have shown a compensatory link between RNA isoform generation and their transcript half-lives, which serves to regulate steady-state mRNA levels under stress conditions (Geisberg et al. 2023). Moreover, the polyA sites from another yeast species *Schizosaccharomyces pombe* show different sequence composition vs. *S. cerevisiae* (Liu et al. 2017). Currently, the polyadenylation complexes in *S. pombe* remain uncharacterized. It remains a question how the polyA sites are determined across yeast species, although experiments showed that 3’UTRs are modular entities that can determine the cleavage and polyadenylation profiles independent of the coding sequences (Lui et al. 2022).

A challenge to characterizing yeast polyA signals is that conventional motif enrichment analysis is hardly applicable to deconvoluting the degenerate motifs and examining the signal differences among genes. The deep learning modeling is well suited to address this question. Specifically, convolutional recurrent neural networks can quantitatively capture dynamic interactions among *cis*-regulatory motifs and resolve the sequence complexity. While a few models have been developed to study human polyA sites (Bogard et al. 2019; Vainberg Slutskin et al. 2019; Chen et al. 2021), they have not been applied to study yeast polyadenylation. Here, we aimed to develop deep learning models to characterize the motif grammar mediating polyadenylation site formation, strength, and cleavage profiles in *S. cerevisiae,* as well as examine the differences in *S. pombe*.

## RESULTS

### Develop a deep learning model named PolyaClassifier to characterize polyA sites in *S. cerevisiae*

To identify genome-wide cleavage and polyadenylation sites in *S. cerevisiae*, we analyzed published 3’ region extraction and deep sequencing (3’READS) data (Table S1) (Hoque et al. 2013; Blair et al. 2016; Liu et al. 2017; Geisberg et al. 2020). After mapping the sequencing reads to reference genome and transcriptome, we selected 15.6 million polyA site supporting (PASS) reads with at least two non-genomic template As to map cleavage sites at the nucleotide resolution and quantify their expression (Figure 1A). As extensive cleavage heterogeneity happens at the yeast 3’-ends (Moqtaderi et al. 2013), we used an iterative approach and selected 38,067 top expressed cleavage sites across 6,094 genes as representatives to develop the deep learning model (see Methods for details). The nucleotide profiles of these sites correspond to the known features: overall enriched for A/U, with an A-rich peak located 10-30 nucleotides (nt) upstream, and U-richness around the cleavage sites (Figure 1B).

**Figure 1.**
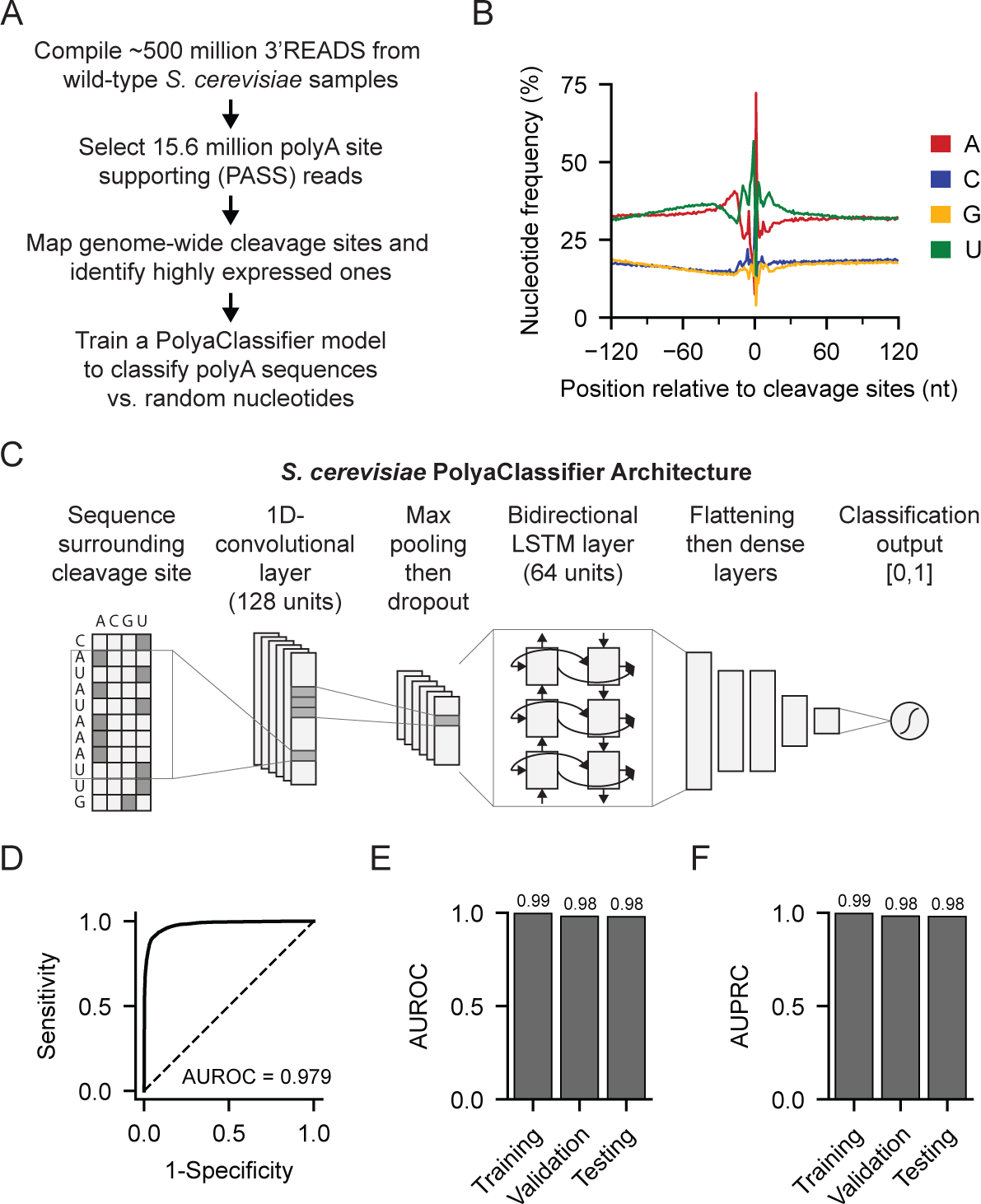
Developing the *S. cerevisiae* PolyaClassifier model. (A) Overview of 3’READS data processing and the model training steps. (B) Distribution of nucleotide frequency surrounding the cleavage sites in *S. cerevisiae*. (C) Architecture of the *S. cerevisiae* PolyaClassifier model, which is described in more detail in Table S2. (D) The ROC curve showing the classification performance of the PolyaClassifier model on the testing set. The AUROC value is shown (N = 7614). (E) The AUROC showing the classification performance of the *S. cerevisiae* PolyaClassifier model on the training, validation, and testing data (N = 60907, 7613, 7614, respectively). (F) Similar to (E), except showing the AUPRC values for the same datasets.

The 240 nt sequences surrounding these cleavage sites were used as positive examples for the model training. Meanwhile, we used an equal number of random genome sequences or shuffled nucleotides as negative examples. We developed a deep neural network model named PolyaClassifier to classify the positive polyA sites vs. negative sequences. The model takes the one-hot embedded sequences as the input, includes one convolutional layer and one bidirectional LSTM layer to capture the 8-mer motif interactions, and performs the site classification (Figure 1C and Table S2; see Methods for detail). Our final model achieved high accuracy in classifying positive and negative sequences. The area under the receiver operating characteristic curve (AUROC) and the area under the precision-recall curve (AUPRC) is 0.99 for the training set, 0.98 for the validation, and 0.98 for the testing (Figure 1D-F).

### Identify *cis*-regulatory motifs and their positional importance in *S. cerevisiae* polyA site formation

Leveraging the PolyaClassifier model, we used a hexamer disruption approach to unbiasedly identify *cis*-regulatory motifs contributing to genome-wide polyA site formation. For a given polyA site sequence, we replaced each hexamer 100 times with random nucleotides and calculated the median log-odds changes (Δlog-odds) in the predicted classification probability. If a hexamer is important for the polyA site formation, we expect that its disruption would cause a decrease in predicted probability. We applied this approach to 34,452 highly expressed polyA sequences (with ≥100 PASS reads and usage level >5%). Then, we calculated the importance score for each hexamer as the sum(Δlog-odds) for each position from −120 to +114 nt around polyA sites. Using a 20-nt scanning window, we identified motifs showing higher sum importance scores than expected and their position-specific importance (Figure 2A, see Methods for detail). Identified motifs belong to known families promoting polyA site formation in *S. cerevisiae*, including the UA-rich motifs (25∼90 nt upstream), A-rich motifs (∼20 nt upstream), and U-rich motifs (around cleavage sites) (Figure 2AB and Table S3). These three motif families showed overall comparable per-site importance scores (Figure 2C).

**Figure 2.**
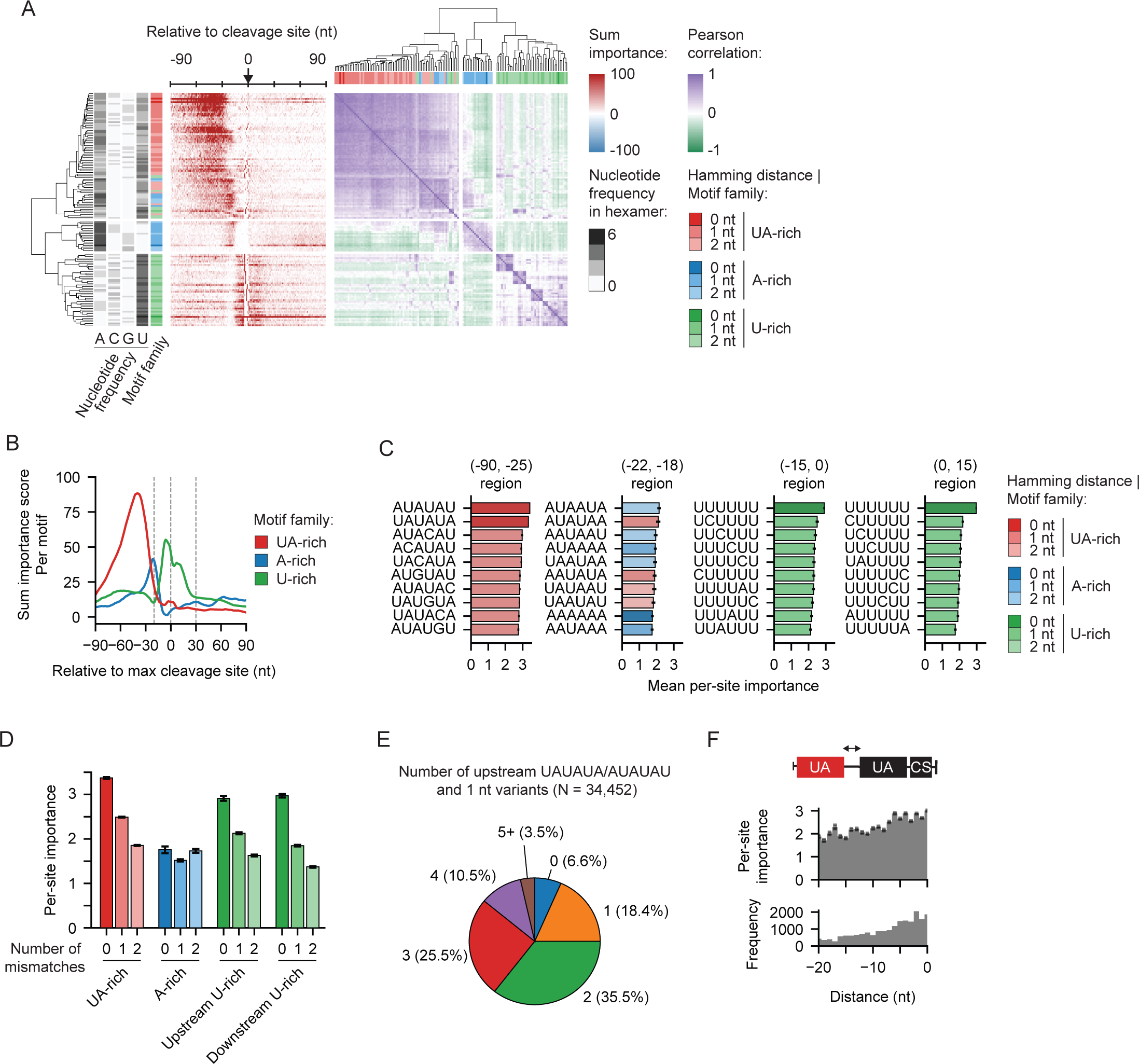
Identifying *cis*-regulatory elements mediating polyA site formation in *S. cerevisiae*. (A) Heatmaps showing the sum classification importance (left) and Pearson correlation between motif importance profiles (right) for *cis*-regulatory elements significantly contributing to polyA site definition (N = 136). Each row represents a hexamer motif and is annotated with the nucleotide content and motif family. Motif families were defined by the Hamming distance between the hexamer and the archetypical motifs UAUAUA/AUAUAU (UA-rich), AAAAAA (A-rich), or UUUUUU (U-rich). (B) The per-motif sum classification importance profiles centered at the maximum cleavage site for the *cis*-regulatory element families. Significant motifs with a Hamming distance ≤2 nt were included for each family. (C) Bar plots showing the per-site importance score of the top 10 motifs in each region surrounding the cleavage site. Bars are colored by the family to which the motif belongs. Data is presented as the mean and the 95% confidence interval (error bar). (D) Bar plot summarizing the mean per-site importance for *cis*-regulatory elements grouped by their motif family and Hamming distance in their region of peak activity. UA-rich motifs occur in the (−90,−25) region, A-rich in the (−22,−18) region, and U-rich in the (−15,0) and (0,15) regions. Data is presented as the mean and the 95% confidence interval (error bar). (E) The fraction of polyA sites (≥100 PASS reads) with non-overlapping upstream efficiency elements, indicated by the presence of AUAUAU/UAUAUA and 1 nt variants (N = 34,452 sites). (F) The per-site importance (top) and frequency (bottom) of upstream UA-rich motifs are grouped by the distance to the last UA-rich motif closest to the cleavage site. Only the two UA-rich motifs located closest to the cleavage sites were included in the analyses (N = 25,841 sites). Data is presented as the mean and the 95% confidence interval (error bar).

Our model quantifies the relative importance of near-cognate motifs. The upstream UA-rich motifs representing the binding of CFIB showed the maximum per-site importance and occurrence at −38 nt (Figure 2B and Figure S1AB). While UAUAUA or AUAUAU showed the maximum importance, their 1-nt variants averaged ∼26% lower importance, and the 2-nt variants showed 45% lower scores (Figure 2CD and Figure S1A). The most important variants include AUACAU, UACAUA, AUGUAU, UAUGUA, which showed 10∼17% lower importance scores than UAUAUA or AUAUAU but similar location enrichment in the upstream region (Figure 2C and Figure S1CD). The majority of polyA sites (75%) have multiple upstream UA-rich elements, with 35.5% containing two and 39.5% having ≥3 (Figure 2E). Then, we examined the optimal distance between the two UA-rich elements closest to the cleavage site. We calculated the per-site importance of the upstream UA-rich element as a function of the distance to the downstream one. The upstream UA-rich elements showed higher importance when the two elements were located within 6 nt (Figure 2F). It is possible that multiple UA-rich elements located close by can increase the efficiency of the CFIB complex binding.

The A-rich motifs located ∼20 nt upstream of the cleavage sites represent the binding sites of CF1A (Figure 2B). Previous reports showed the presentative motifs are AAAAAA or AAUAAA (Mónica et al. 2011). However, AUAAUA, AUAUAA, AAUAAU, and AUAAAA show the highest per-site importance score in the region (Figure 2C). The presence of 1 or 2 Us in AAAAAA did not decrease the motif importance (Figure 2D and Figure S1A). Some motif variants, such as AUAUAA or UAAAUA, are likely to function as the binding sites of both CFIB (UA-rich variants) and CF1A (A-rich variants), considering the distribution of their occurrences and per-site importance scores (Figure S1B).

The U-rich elements bound by the CPF are located within 20 nt both upstream and downstream of the cleavage site (Figure 2AB and Figure S1A). On average, 1-nt variants of UUUUUU decreased the per-site importance score by 32%, while 2-nt variants decreased the score by 49% (Figure 2CD and Figure S1A). The variants showing top importance scores contained 1 nt of C or A within UUUUUU (Figure 2C). Altogether, we used the PolyaClassifier model and quantitatively revealed the relative importance of degenerate motifs in promoting *S. cerevisiae* polyA site formation.

### Examine molecular mechanisms mediating the cleavage heterogeneity in *S. cerevisiae*

PolyA sites in *S. cerevisiae* show different levels of cleavage heterogeneity. Some sites show high cleavage heterogeneity spanning > 30 nt, while for others, the cleavage sites are relatively precise (Figure 3A and Figure S2A). We developed an entropy score to quantify the extent of cleavage heterogeneity of a polyA site. For each cleavage site region, we selected the representative site showing the highest number of PASS reads and measured the uniformness of the read distribution in the surrounding +/−25 nt region (See Methods for details). A higher entropy value indicates a higher number of unique 3’-ends generated during mRNA processing (Figure 3A and Figure S2A).

**Figure 3.**
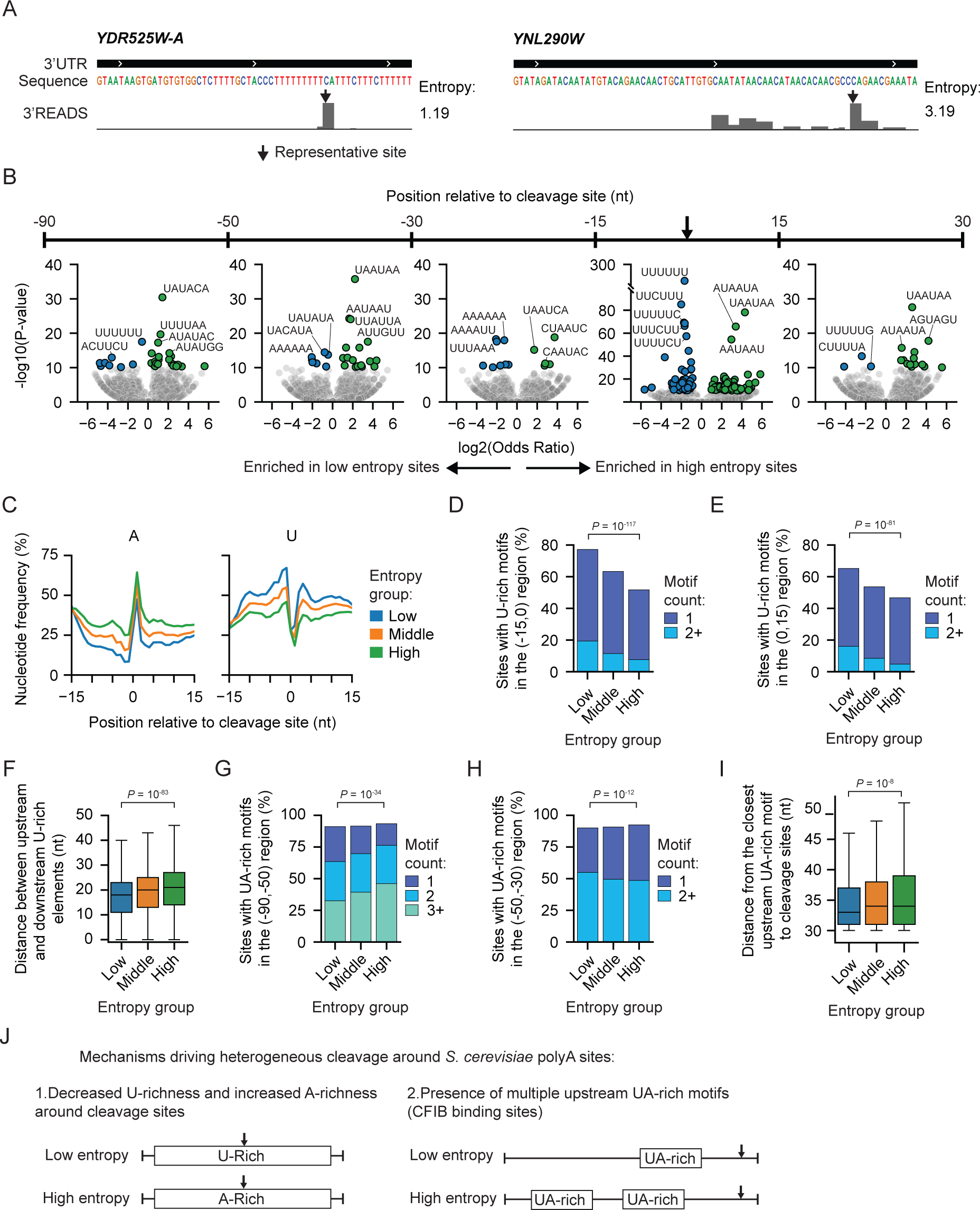
PolyA site heterogeneity is driven by cleavage site composition and upstream efficiency elements. (A) Examples of the PASS read distribution surrounding a low entropy site (left) and a high entropy site (right). The entropy values are shown. (B) Motif enrichment analysis comparing the low (bottom 20%) vs. high (top 20%) entropy polyA site groups. Five different polyA site regions were analyzed as indicated in the figure. The x-axis indicates the degree of enrichment as measured by the log2(odds ratio). The y-axis shows the - log10(Chi-squared test *P*-value) for the motifs enriched in the high entropy group and log10(Chi-squared test *P*-value) for ones enriched in the low entropy sites. (C) The nucleotide distribution flanking the maximum cleavage sites showing the fraction of As (left) and Us (right). (D) The fraction of sites in each entropy group containing one or more U-rich motifs in the (−15,0) region immediately upstream of the maximum cleavage site. U-rich motifs with up to 2 nt mismatches from UUUUUU were included. The *P*-value from the Chi-squared test for independence across entropy groups is shown. (E) Similar to (D), except showing the analyses in the (0,15) region immediately downstream of the maximum cleavage site. (F) Boxplots showing the distance between the closest upstream and downstream flanking U-rich elements. The *P*-value is the result of a Wilcoxon rank-sum test comparing the low vs. high group. (G) The fraction of sites in each entropy group containing one or more UA-rich motifs in the (−90,−50) region upstream of the maximum cleavage site. UA-rich motifs with up to 2 nt mismatches from AUAUAU or UAUAUA were included. The *P*-value from the Chi-squared test for independence across entropy groups is shown. (H) Similar to (G), except showing the fraction of sites with UA-rich motifs in the (−50,−30) region upstream of the maximum cleavage site. (I) Boxplots showing the distance from the closest UA-rich motif in the (−90,−30) region to the cleavage site (N = 3025 low, 9105 middle, and 3028 high entropy sites). (J) Schematic depicting the mechanisms identified from this analysis that may lead to heterogeneous cleavage in *S. cerevisiae*.

We performed *cis*-element enrichment analyses comparing polyA site sequences showing high vs. low cleavage heterogeneity (comparing the sites with top 20% vs. low 20% entropy scores; Figure 3B). The most significant difference was found in the −15 to +15 nt around the polyA sites. While the precise cleavage sites showed enrichment of U-rich elements (e.g. UUUUUU, UUCUUU, and UUUUUC) in the region, the high heterogeneous sites were enriched with A-rich elements (e.g. UAAUAA, AUAAUA, and AAUAAU) (Figure 3B; Chi-squared test *P*-value < 10^−40^). Considering the frequency of single nucleotides, the high heterogeneous cleavage sites show lower U-richness and higher A-richness in the sequence surrounding the maximum cleavage site (Figure 3C and Figure S2B-D). Next, we examined the occurrence of U-rich hexamers defined in Figure 2 (binding sites of the CPF complex). High heterogeneity sites are less likely to have U-rich hexamers flanking the cleavage site vs. the low heterogeneity group (Δfraction of sites = 20%) (Figure 3DE). Some sites in the high heterogeneous group have both upstream and downstream U-rich elements, but the two elements tend to be located farther apart than in the low heterogeneous group (Figure 3F). Altogether, the data indicated that polyA sites showing high heterogeneous cleavage are devoid of U-rich elements around cleavage sites which can lead to the weaker binding of the CPF complex.

Another major difference between high and low heterogeneity cleavage sites was the frequency of UA-rich elements located 30 ∼ 90 nt upstream (the binding sites of CF1B). The motif enrichment analyses showed the UA-rich motifs (e.g., UAUACA and AUAUAC) occur more frequently in the upstream 50-90 nt region but are found lower frequency in the 30-50 nt region in the high heterogeneity sites (Figure 3B and 3GH). Considering the closest upstream AU-rich element, these motifs were farther away from the cleavage site in the high heterogeneity group (Figure 3I). Taken together, our above analyses showed that the heterogeneous cleavage sites in *S. cerevisiae* tend to show the depletion of U-rich elements around polyA sites and have a higher occurrence of upstream UA-rich motifs (Figure 3J).

### Develop a deep learning model named PolyaCleavage to capture the cleavage heterogeneity of polyA sites based on primary sequences

We reasoned that if the polyA site sequences determine the extent of cleavage heterogeneity, we should be able to capture that regulation by deep learning modeling. To this end, using the 240-nt sequences centered at the maximum cleavage sites as the input, we developed another deep learning model named PolyaCleavage to predict the cleavage probability distribution in the middle 50 nt region (Figure 4A and Table S2; see Methods for details). To evaluate the performance of our algorithm, we calculated the entropy values measuring the cleavage heterogeneity based on the PolyaCleavage prediction. The predicted entropy values were correlated with those calculated using the observed 3’READS data (Figure 4B). Additionally, the weighted mean cleavage positions were well correlated between the prediction and the observed 3’READS, with a correlation coefficient of 0.82 for the testing set (Figure S3A). These data indicated that our PolyaCleavage model quantitatively captured the cleavage heterogeneity based on the primary sequences.

**Figure 4.**
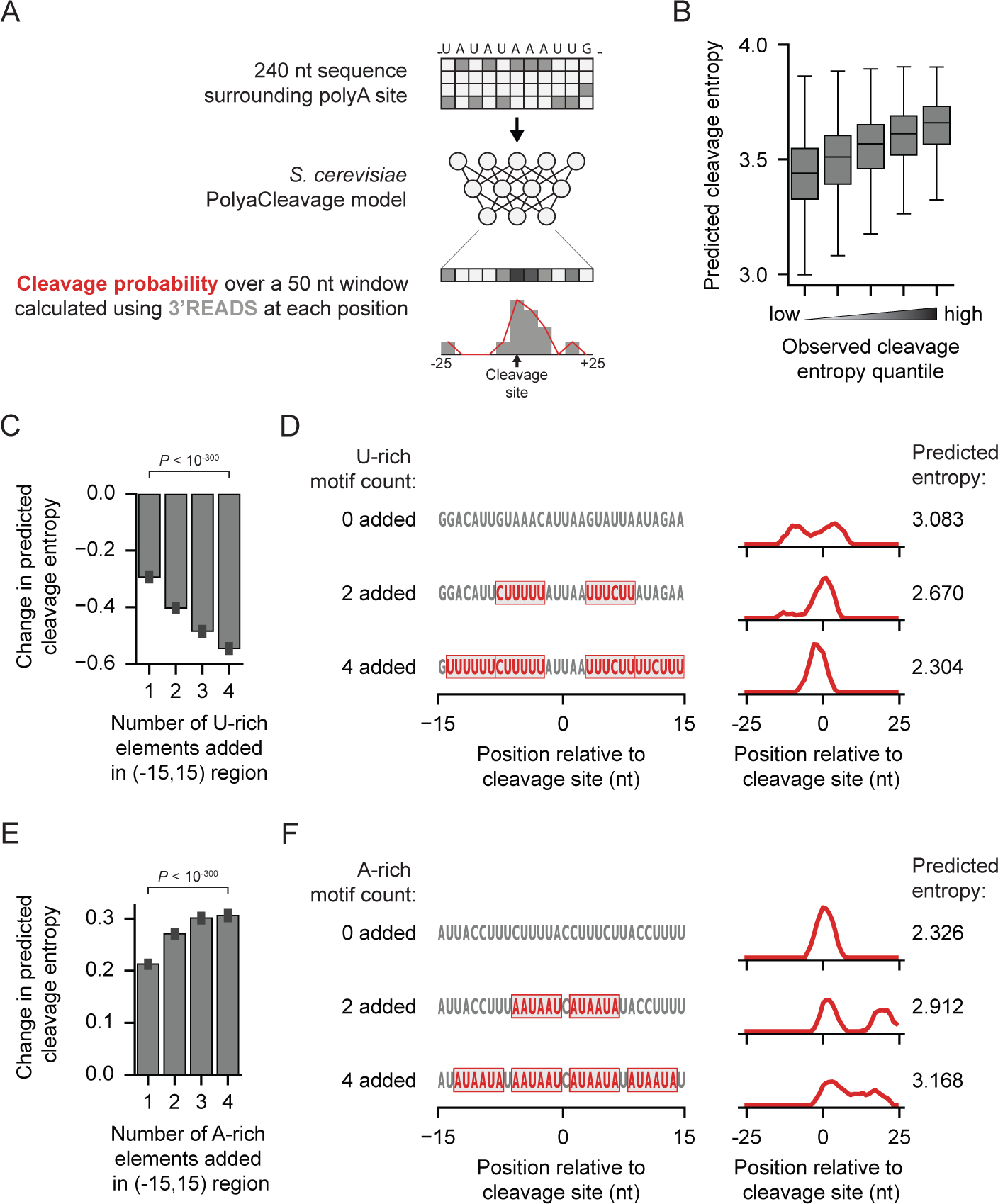
Cleavage heterogeneity can be modulated by changing the polyA site surrounding *cis*-regulatory elements. (A) Workflow describing our design of *S. cerevisiae* PolyaCleavage model. (B) Boxplots showing the increase in predicted cleavage entropy as the observed cleavage entropy measured by 3’READS increases. PolyA sites from the holdout testing set supported by at least 50 PASS reads were included. These sites were split into 5 evenly sized groups (N = 6538 sites per group). (C) The change in predicted cleavage entropy as non-overlapping U-rich elements are sequentially introduced in the (−15,+15) region surrounding the maximum cleavage site. The data are presented as the mean and the 95% confidence interval (error bar). The *P*-value from the Wilcoxon rank-sum test comparing the addition of 1 U-rich element to 4 U-rich elements is shown. (D) An example showing the effect of introducing U-rich elements around the cleavage site in a high entropy polyA site from *YGR238C*. The predicted entropy values are shown. (E) The change in predicted cleavage entropy as non-overlapping A-rich elements are sequentially introduced in the (−15,+15) region immediately surrounding the maximum cleavage site. The data are presented as the mean and the 95% confidence interval (error bar). The *P*-value from the Wilcoxon rank-sum test comparing the addition of 1 A-rich element to 4 A-rich elements is shown. (F) Similar to (D), showing the effect of introducing A-rich elements around the cleavage site in a low entropy polyA site from *YDL202W*.

Next, we leveraged the PolyaCleavage model to study the genomic parameters we identified above in regulating cleavage heterogeneity. We first examined the roles of U-rich motifs around the cleavage sites. Adding U-rich motifs to the +/−15 nt region around highly heterogeneous polyA sites induced more precise cleavage as predicted by the PolyaCleavage model (Figure 4CD and Figure S3B). On the other hand, replacing the U-rich motifs with the A-rich elements around the sites was predicted to increase cleavage heterogeneity (Figure 4EF and Figure S3C). The predicted change in entropy was correlated with the number of U-rich or A-rich elements introduced. We also examined the effects of upstream CF1B binding sites by reducing the number of UA-rich elements in −90 nt to −30 nt regions. We observed more precise cleavage as we reduced the number of UA-rich elements (Figure S3DE). Altogether, by developing the PolyaCleavage model, we showed that the sequence surrounding polyA sites determines the degree of cleavage heterogeneity in *S. cerevisiae* and confirmed the regulatory roles of identified factors.

### A deep learning model named PolyaStrength to quantify *S. cerevisiae* polyA site strength

To quantify the polyA site expression levels, we used an iterative approach and grouped nearby cleavage sites (within 20 nt) into clusters to account for the heterogeneous cleavage (Table S4; see Methods for details). We used the 20-nt window because we showed that an A-rich motif located 20 nt upstream was required for downstream polyadenylation (Figure 2B). The expression level of a polyA site (defined by the cluster) was calculated using the summed PASS read count in the region. For genes with multiple polyA sites in the 3’UTR, we calculated the usage level of each site as the ratio between the number of supporting reads vs. the sum of reads supporting the top two expressed sites.

A major regulator of polyA site usage levels is the efficiency of *cis*-regulatory elements in recruiting the polyadenylation factors. Next, we developed a deep learning model named PolyaStrength using the 240 nt sequences centered at the maximum cleavage sites as the input to predict the site usage levels on the log-odds scale (Figure 5A and Table S2; see Methods for details). Our PolyaStrength score is correlated with 3’UTR polyA site usage levels (Figure 5B) and effectively classified the highly vs. lowly used polyA sites (>8-fold expression level differences) with the AUROC and AUPRC values >0.92 for the training, validation, and testing sets (Figure 5C and Figure S4AB).

**Figure 5.**
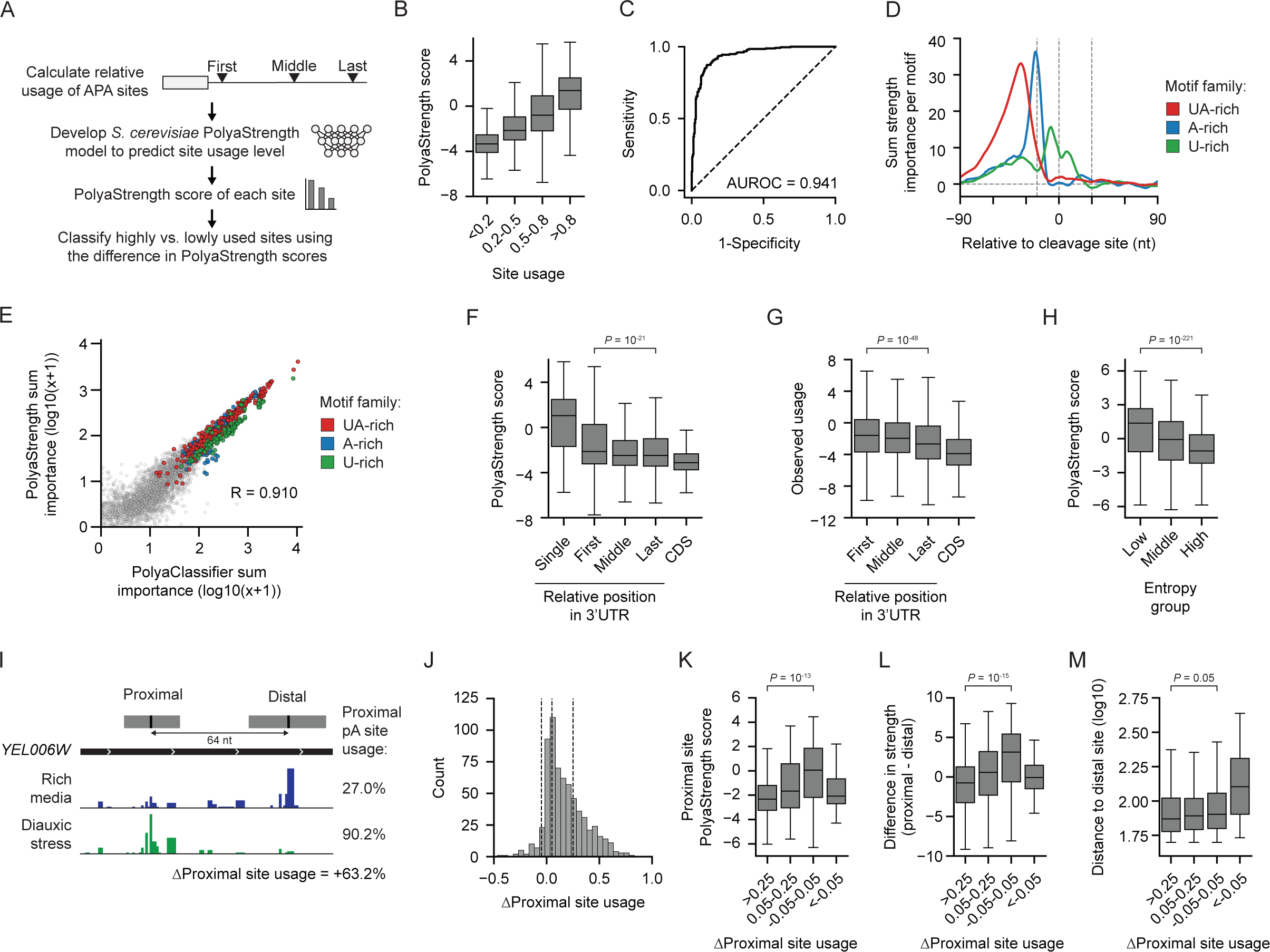
Development of the *S. cerevisiae* PolyaStrength model. (A) Diagram depicting how the *S. cerevisiae* PolyaStrength model was trained on the relative usage scores for APA sites in the 3’UTR. (B) Boxplot showing the predicted PolyaStrength scores grouped by observed site usage levels. (C) The ROC curve showing the performance of the *S. cerevisiae* PolyaStrength model to distinguish lowly vs. highly used 3’UTR APA sites from the holdout testing set (N = 495 pairs of sites showing ≥8-fold differential usage levels). (D) The per-motif sum importance scores around the cleavage sites for indicated *cis*-regulatory element families. All significant motifs with a Hamming distance ≤2 nt from the archetypical motifs UAUAUA/AUAUAU, AAAAAA, and UUUUUU were included. (E) Scatter plot showing the correlation between the motif importance scores for the *S. cerevisiae* PolyaClassifier and PolyaStrength models. The peak sum importance score in any 20 nt window is used and the Pearson correlation is shown. (F) The predicted PolyaStrength scores for polyA site types are based on their relative position. The *P*-value from the Wilcoxon rank-sum test comparing the first vs. last 3’UTR-APA sites is shown. (G) The observed usage levels of coding region and 3’UTR-APA sites. See Methods for details about how the site usage is calculated. The *P*-value from the Wilcoxon rank-sum test comparing first vs. last 3’UTR-APA sites is shown. (H) The predicted PolyaStrength scores for low, middle, and high entropy polyA sites from Figure 3. The *P*-value using the Wilcoxon rank-sum test comparing the low vs. high entropy groups is shown. (I) An example gene *YEL006W* shows increased usage of proximal polyA site under diauxic stress. The proximal and distal polyA site clusters are shown above, with the distance between them noted. The usage levels of the proximal site are shown. (J) The distribution of delta usage levels for proximal sites shows significantly different usage under diauxic stress compared to rich media conditions (N = 660). The vertical dashed lines show the group of differentially used sites based on their usage level changes. (K) The PolyaStrength scores of the proximal sites grouped by the Δusage. The *P*-value from the Wilcoxon rank-sum test comparing the high Δusage group (>0.25) and the low Δusage group (−0.05 – 0.05) is shown. (L) Similar to (K), except showing the strength differences between the proximal site and its paired distal site. (M) Similar to (K), except showing the distance between the paired proximal and distal sites.

Using the PolyaStrength model, we used the hexamer disruption approach as we used above to identify *cis*-regulatory elements contributing to the site strength. Similar motifs were identified compared to those from the PolyaClassifier model, including the UA-rich elements (−90 ∼ −30 nt), A-rich elements (∼ −20 nt), and U-rich elements around the cleavage site (Figure 5D and Table S5). The sum importance scores of the motifs were highly correlated between the PolyaStrength and PolyaClassifer models, indicating that similar motifs promote polyA site strength and formation (Figure 5E).

We grouped the polyA sites based on their relative genomic locations in 3’UTRs: single sites in non-APA genes, as well as first, middle, or last sites in APA genes (Table S4). The single polyA sites were stronger than other types, and the first polyA sites were generally stronger and showed higher usage levels than other APA sites (Figure 5FG). The sites located in coding regions tend to be weaker and have lower usage levels than those in 3’UTRs (Figure 5FG). PolyA sites showing high cleavage heterogeneity were also weaker than those with precise cleavage (Figure 5H).

Next, we examined the APA regulation under the diauxic stress. A previous study showed a global increase in proximal site usage under stress conditions due to slower elongation of RNA polymerase II (Geisberg et al. 2020). However, the proximal sites showed differential usage levels (Figure 5IJ and Figure S4C-E). The underlying reasons remain uncharacterized. We grouped the polyA sites based on their usage level fold changes (Figure 5J) and observed the correlation between site strength and the APA regulation. The proximal polyA sites that showed higher usage increase under diauxic stress were weaker considering their absolute and relative strengths (Figure 5KL). Consistent with the previous report (Geisberg et al. 2020), a minor fraction of proximal polyA sites (5%) showed decreased usage. These sites were weaker and located further away from the corresponding distal sites (Figure 5K-M). Altogether, we developed the PolyaStrength model to quantify the polyA site strengths, which can be useful for dissecting the molecular mechanisms underlying APA regulation during biological processes.

### Examine the motif composition of polyA sites in *S. pombe* using deep learning

*S. pombe* is another commonly used yeast model species. Unlike *S. cerevisiae,* the configuration of the polyadenylation machinery in *S. pombe* remains mostly uncharacterized. Previous 3’READS profiling and analyses showed that nucleotide profiles surrounding *S. pombe* polyA sites differ from *S. cerevisiae* or mammals (Liu et al. 2017). Overall, the polyA region is highly AU-rich, with an A-rich peak located at 20 nt upstream and U-richness around the cleavage sites (Figure 6A and Table S6). Using a similar approach to our analysis of *S. cerevisiae*, we built a PolyaClassifer model to distinguish *S. pombe* polyA sequences vs. random nucleotides (Figure 6B). Our model classified the sites well, with AUROC and AUPROC values ≥0.95 for training, validation, and testing sets (Figure 6C-E).

**Figure 6.**
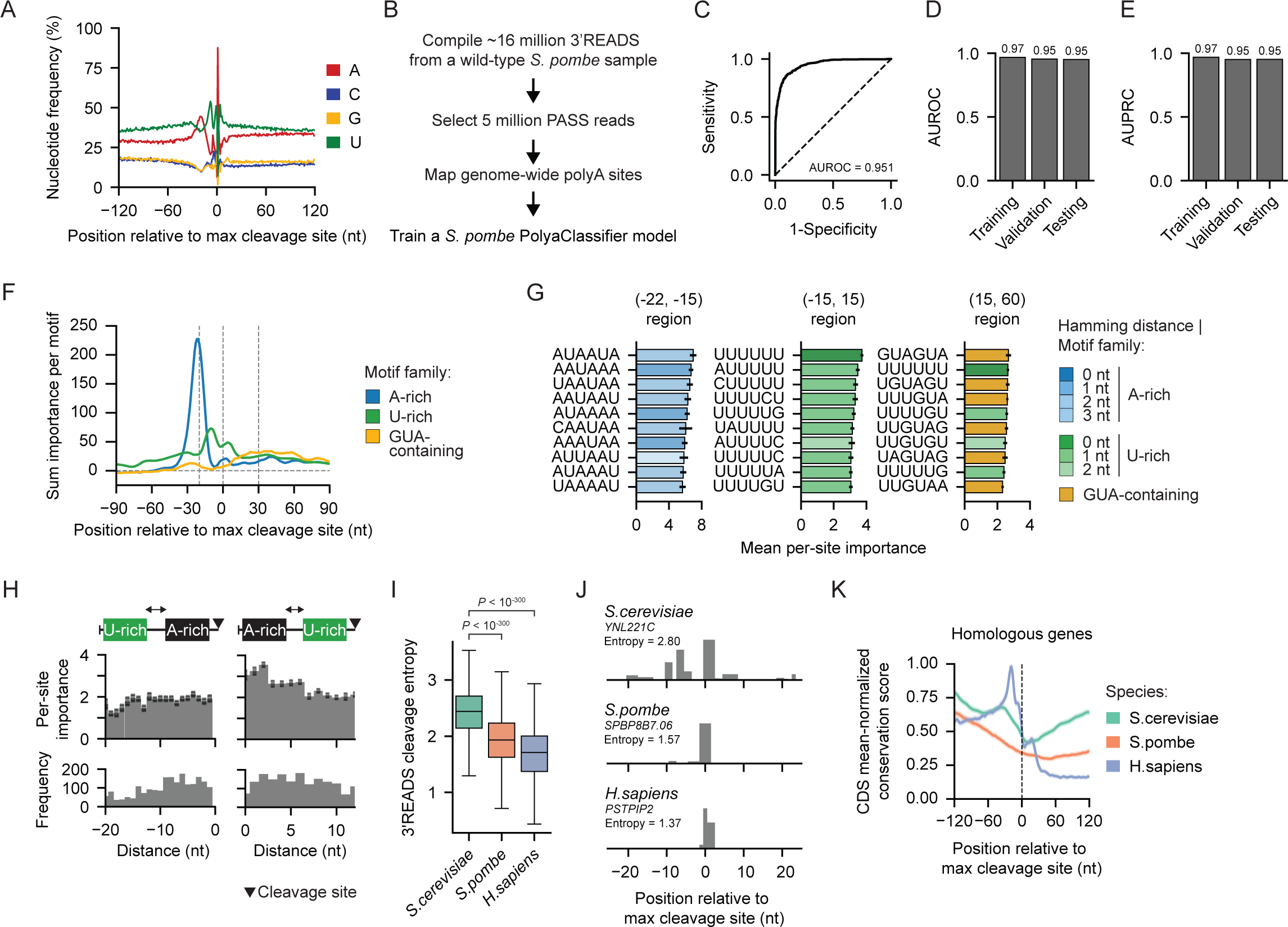
The *S.pombe* PolyaClassifier model reveals divergent motifs mediating polyA site definition. (A) The nucleotide profiles around the *S. pombe* polyA sites. (B) Overview of the data processing workflow to train the *S. pome* PolyaClassifier model. (C) The ROC curve showing the model performance for the holdout testing set. (D) The AUROC values for the training, validation, and testing sets (N = 9851, 1231, 1232, respectively). (E) Similar to (D), except showing the AUPRC values. (F) The per-motif sum classification importance profiles centered at the maximum cleavage site. (G) Bar plots showing the per-site importance of the top 10 motifs in each region surrounding the maximum cleavage site. Bars are colored by the family to which the motif belongs. Data is presented as the mean and the 95% confidence interval (error bar). (H) The per-site importance (top) and frequency (bottom) for U-rich motifs are grouped by their distance upstream (left) or downstream of the A-rich motif (right). Data is presented as the mean and the 95% confidence interval (error bar). (I) The distribution of 3’READS cleavage entropy values for the top polyA site in each gene homologous across *S. cerevisiae, S. pombe,* and *H. sapiens* (N = 6546, 6373, and 8170, respectively). The *P*-values from the Wilcoxon rank-sum tests comparing *S. cerevisiae* to *S. pombe* and *H. sapiens* are shown. (J) An example of the PASS read distribution +/−25 nt surrounding the top-expressed polyA site in homologous genes: *S. cerevisiae* gene *YNL221C, S. pombe* gene *SPBP6B7.06, and H. sapiens* gene *PSTPIP2*. The cleavage entropy value calculated from the 3’READS for each species is shown. (K) A metaplot showing the conservation score for the +/−120 nt surrounding the top expressed polyA sites in all homologous genes across the three species. The conservation score was normalized to the mean score in coding regions. For APA genes, we only selected the top expressed site in the analyses. The data is shown as the mean with the shaded region indicating the 95% confidence interval.

The hexamer disruption analyses revealed the motifs that contributed to polyA site formation, including the A-rich motifs located −20 nt upstream, U-rich motifs surrounding the cleavage sites, and GUA-containing motifs +15 to +60 nt downstream (Figure 6F, Figure S5A, and Table S7). The upstream A-rich motifs can be quite degenerate and the ones with the highest importance scores tend to contain one or two Us, such as AUAAUA, AAUAAAA, and UAAUAA (Figure 6G). Considering the per-site importance, the A-rich motifs showed an overall ∼2-fold higher score than the U-rich and GUA-containing elements (Figure 6G and Figure S5B). The U-rich motifs were found across a broad region from 30 nt upstream to 10 nt downstream of the cleavage sites (Figure S5B). The U-rich motifs showed the maximum importance when they were immediately downstream of the A-rich motif (<6 nt) (Figure 6H). The GUA-containing motifs were more important in the context of upstream Us and downstream G (Figure 6G and Figure S5C). Our results revealed the motif configuration of *S. pombe* polyA sites which is distinct from *S. cerevisiae*.

Using our entropy calculation method, we quantified the cleavage heterogeneity of *S. pombe* polyA sites. Overall, the cleavage heterogeneity was much lower than in *S. cerevisiae* and we did not observe large cleavage “zones”, although the overall heterogeneity was larger than in homologous human genes (Figure 6IJ and Figure S5D). Next, we examined the evolution of polyA site sequences across model species using the PhastCons scores calculated based on the multiple genome alignments (Siepel et al. 2005; Rhind et al. 2011). We examined the sequence conservation of *S. pombe* polyA sequences using the alignments across 4 *Schizosaccharomyces* yeasts, the conservation of *S. cerevisiae* sites across 7 *Saccharomyces* yeasts, and the human site conservation across 100 vertebrates. For human polyA sites, we observed high sequence conservation upstream of the cleavage site which peaked at the PAS region (AAUAAA and variants located at −20 nt upstream) (Figure 6K). However, we did not observe the conservation peak at the A-rich elements (−20 nt) for *S. cerevisiae* or *S. pombe* polyA sites (Figure 6K and Figure S5E). Using the coding regions as the control, in yeast, the conservation levels gradually decreased toward the polyA sites (Figure 6K and Figure S5E). The trend was consistent when we analyzed different polyA site types in the 3’UTRs (i.e., single, first, middle, or last) (Figure S5F). The results could be partly due to the degenerate nature of the yeast polyA signals but also indicated that the polyA regions of yeast species are under positive selection.

## DISCUSSION

The yeast species have been valuable model systems to dissect basic molecular mechanisms mediating steps of gene expression. While the transcription initiation and elongation have been well characterized using these organisms, the regulation of cleavage and polyadenylation has been understudied. Only recently, studies used deep sequencing approaches to map genome-wide polyA sites and examined the functional roles of different 3’-end RNA isoforms in *S. cerevisiae*. An unexpected finding was the extensive heterogeneous cleavage for some mRNA genes. Although the conventional model showed important motifs contributing to polyA site formation, it is not sufficient to explain the cleavage heterogeneity and signal differences among genes. It has been puzzling how the 3’-ends of mRNAs are determined in *S. cerevisiae.* Here, we developed the first deep learning models to examine motif grammars mediating polyA site formation, heterogeneity, and strength in *S. cerevisiae*.

Previous molecular experiments of several polyA sites indicated that the *cis*-regulatory elements mediating *S. cerevisiae* polyadenylation can be quite degenerate. However, no study has been carried out to systematically examine the relative importance of signal variants. Our deep learning models PolyaClassifer and PolyaStrength provided unprecedented insights into the degenerate nature of the *cis*-regulatory motifs in *S. cerevisiae*. Using a hexamer disruption approach, we unbiasedly identified the positional importance of motifs mediating *S. cerevisiae* polyA site formation and strength. The significant motifs belong to the previously defined families: the CFIB bind site (the upstream UA-rich elements), the CF1A binding sites (A-rich motifs), and the CPF binding sites (the U-rich elements around cleavage sites).

Our analyses provided quantitative measurements of the near-cognates’ strengths. For the CFIB binding site, replacing one nucleotide of the consensus motif UAUAUA or AUAUAU sometimes decreases their strength by only ∼15%. An unexpected finding is that most polyA sites contain multiple upstream UA-rich elements, suggesting that the CF1B complex or some subunit protein(s) could form dimers like the mammalian CstF complex. And for the CF1A binding, adding one or two Us in AAAAAA does not change motif strengths and sometimes can even make the motif stronger. The location of the A-rich element is like the PAS motif in mammals (AAUAAA or variants), which is 20 nt upstream of polyA sites. However, the CF1A binding site can be much more degenerate than mammalian PAS, of which AAUAAA is much stronger than other variants. This degenerate nature of motifs in *S. cerevisiae* can create many possible polyA site combinations.

We used an entropy score to quantify the cleavage heterogeneity of polyA sites. Our comparative analyses revealed sequence differences between sites showing high vs. low cleavage heterogeneity. First, the depletion of U-rich elements and enrichment of A-rich motifs surrounding polyA sites promoted heterogeneous cleavage. This could reduce the binding affinity of the CPF complex. A possible model is that for the high heterogeneity cleavage sites, the upstream binding sites of CF1A and CF1B are sufficient to recruit the cleavage and polyadenylation complexes, including the CPF complex. However, CPF binding is not stable due to the lack of U-rich elements, causing cleavage to occur across multiple A-rich sites. In fact, previous experiments showed that the required polyadenylation signals in *S. cerevisiae* include only the CF1A and CF1B binding sites and the polyA sequence itself (i.e., A-rich sequence at the cleavage site) (Guo and Sherman 1996). Later studies showed that the U-rich elements flanking polyA sites are bound by the CPF complex and enhance the polyadenylation activities (Dichtl and Keller 2001). These experimental results are in line with our findings. Second, the increased numbers of CF1B binding sites (AU-rich elements) can lead to different conformation positions of the polyadenylation machinery and increased cleavage heterogeneity downstream.

By developing the PolyaCleavage model, we confirmed that the primary polyA sequences encode the cleavage heterogeneity. We could capture the dynamic changes of cleavage heterogeneity by mutating key motifs. Adding U-rich motifs surrounding polyA sites led to more precise cleavage, while adding A-rich motifs increased heterogeneity. We also confirmed the effects of the number of upstream CF1B binding sites. These mechanisms, in combination with the degenerate nature of yeast *cis*-elements, drive the high number of cleavage and polyadenylation sites utilized in *S. cerevisiae* genes.

We also developed a PolyaStrength model to measure the polyA site strength, which is determined by similar motifs as described above for site formation. The single polyA sites showed the highest strength. For APA genes, the proximal polyA site tends to be stronger than the downstream ones. This differs from mammalian genes, where the distal polyA sites are generally stronger (Hoque et al. 2013; Wang et al. 2018). The sites showing highly heterogeneous cleavage tend to be weaker, which is in line with the fact that they are generally depleted of U-rich motifs around the polyA site. Our PolyaStrength model is useful for future studies of APA regulation during biological processes. Here, we showed that under diauxic stress, the differential usage levels of polyA sites are correlated with the site strength and tandem site distances.

Finally, we examined the polyA site composition in *S. pombe* and found their site configuration is divergent from *S. cerevisiae* and mammals. Currently, the cleavage and polyadenylation complex in *S. pombe* remains uncharacterized. For some features, *S. pombe* sites resemble mammalian sites. The cleavage activity is more precise than in *S. cerevisiae*, and we rarely observe a large heterogeneous cleavage zone*. S. pombe* sites also require an A-rich motif located 20 nt upstream, which showed much higher importance scores than other motifs (i.e., U-rich or UGA-containing motifs) comparable to the PAS in mammals. However, the A-rich motif can be quite degenerate, with tens of elements showing comparable strength.

Taken together, our study provided unprecedented insights into polyA site formation and cleavage heterogeneity in *S. cerevisiae* and *S. pombe*. The deep learning tools we developed can facilitate the study of the functional roles of APA regulation during biological processes. Our results also underscored the need for future molecular experiments to characterize the structures of the polyadenylation machinery in yeast species.

## METHODS

### Identifying genomic polyA sites analyzing 3’READS data

We used 3’READS data to comprehensively identify polyA sites across the *Saccharomyces cerevisiae* and *Schizosaccharomyces pombe* genomes (Hoque et al. 2013; Blair et al. 2016; Liu et al. 2017; Geisberg et al. 2020). We compiled data from 7 *S. cerevisiae* and 1 *S. pombe* samples for wild-type yeast cultured under rich media conditions (Table S1). We analyzed the 3’READS data as the following. The leading Ts were trimmed from each read and we logged the number of Ts for downstream use. The trimmed reads were aligned to the transcriptome and then the genome using TopHat v2.1.0 (Kim et al. 2013). For *S. cerevisiae*, we used the Ensembl reference genome version R64-1-1 and the gene annotation curated by the Struhl Laboratory. For *S. pombe*, we used the Ensembl reference genome assembly ASM294v2 and the RefSeq gene annotation from the UCSC Genome Browser.

The uniquely mapped reads with ≥2 non-genomic As named the polyA site supporting (PASS) reads were used for the downstream analyses identifying polyA sites. Our last alignment site is located at a non-A position. However, the cleavage tends to happen in A-rich regions (Dichtl and Keller 2001). For some sites, there are genomic As after the non-A site and we cannot distinguish where the cleavage happens among the positions. We evenly distributed the PASS reads from the first non-A to the last As. To assign cleavage sites to their respective genes, we extended 3’-ends of annotated genes up to 500 nt or until the transcription start site of the downstream gene on the same strand without overlap.

### Developing the PolyaClassifier model to classify polyA site sequences vs. random nucleotides in *S. cerevisiae* and *S. pombe*

#### Input dataset

To train the model, we identified high-confidence cleavage sites. For *S. cerevisiae*, we required that a cleavage site be supported by ≥10 PASS reads and ≥2% of reads from the maximum expressed site of a gene. We had much fewer reads for *S. pombe* and required that a cleavage site should contain ≥5 PASS reads. In total, we identified 150,232 individual cleavage sites in *S. cerevisiae* and 42,730 in *S. pombe*. We used an iterative approach to sample a subset of cleavage sites as representatives to train the model. We first sampled the highest expressed cleavage site from each gene. Next, we selected the positive sites located ≥5 nt away from the previously sampled sites and picked the next highest expressed site. Using this iterative method, we sampled 38,067 representative cleavage sites in *S. cerevisiae* and 6,157 sites in *S. pombe.* The 240 nt sequences around these cleavage sites (+/−120 nt) were used as positives to train the model.

We generated an equal number of negative control sequences (240 nt) from the following three types: (1) shuffled sequences from the positive cleavage sites maintaining the nucleotide frequencies; (2) random genome sequences without the cleavage sites. For the model input, each sequence was one-hot encoded to a 4×240 matrix where the rows represent the nucleotides A, C, G, and U and the columns represent the position along the sequence. At each position, the row corresponding to the nucleotide present in the genomic sequence is marked with a 1, and all others are marked with a 0. This numerical encoding was fed into the model as input, paired with the binary classification label of 1 if the site is positive and 0 if it is negative.

#### Model architecture and training

The PolyaClassifier model is constructed using a 1D convolutional layer followed by max pooling and dropout layers. It then contains a bidirectional long short-term memory (LSTM) layer followed by a dropout layer. The output of these layers is then flattened and fed into a series of 4 dense layers. The final dense layer uses a sigmoid activation to predict the classification probability of each site. The 1D convolutional layer used a filter size of 8 nt and a stride of 1. This combination of layers extracts informative sequence motifs and their spatial position within the sequence. Further details about the model architecture are in Table S2.

We partitioned the input datasets (positive and negative sequences) into 80% training, 10% validation, and 10% holdout testing sets. We then trained the PolyaClassifier models using a batch size of 100 with a learning rate of 0.001 and the Adam optimizer with Nesterov momentum. The model was trained to minimize the binary-cross entropy loss function, implemented out of the box by Tensorflow 2:

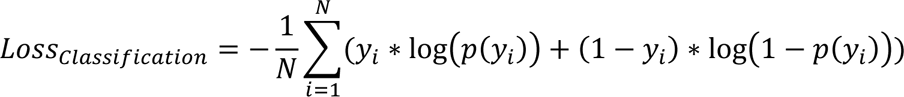

where *y*_*i*_ is the classification label for the sequence *i* (1 for positive site and 0 for negative sequence) and *p*(*y*_*i*_) is the predicted probability of a sequence *i* being a positive polyA site. After each training epoch, the loss and accuracy were tracked for the training and validation splits. The models reached the minimum loss for the validation split and training was stopped after 25 epochs for *S. cerevisiae* and 24 epochs for *S. pombe*.

#### Performance evaluation

To evaluate the performance of the PolyaClassifier model, we calculated the AUROC and AUPRC for the training, validation, and holdout test set using the Python library scikit-learn.

### Characterizing *cis*-regulatory elements modulating polyA site formation

We used a systematic hexamer disruption approach to identify the *cis*-regulatory elements that contribute to the PolyaClassifier predictions and therefore are important in defining polyA site. We replaced each hexamer in the sequence surrounding a polyA site with randomized nucleotides 100 times without altering the rest of the sequence. We then calculated the median change in model predictions for each hexamer occurrence using the logit-transformed PolyaClassifier probability (log-odds).

We employed this technique to discover the motifs defining polyA sites genome-wide in *S. cerevisiae* and *S. pombe*. We carried out motif disruption for 34,452 well-expressed cleavage sites in *S. cerevisiae* (≥100 reads and ≥5% usage) and 7,872 in *S. pombe* (≥50 reads and ≥5% usage). For each position from −120 nt to +120 nt surrounding the maximum cleavage site, we quantified the sum importance score for each hexamer by taking the sum of the disruption effect. We also calculated the per-site importance dividing the sum importance score by the motif frequency. Doing this for each position surrounding the maximum cleavage site created an importance profile showing the position-specific effects of each hexamer.

Next, we characterized the *cis*-regulatory elements that significantly contributed to site definition using a two-step filtering procedure. Previous work has shown that yeast polyadenylation *cis*-regulatory elements are optimally located within 40 nt of the cleavage site (Tian and Graber 2011). First, we shuffled the sum importance scores from the regions outside (−40,40) and constructed a background distribution of 20 nt window scores. We used the 99^th^ percentile score from the distribution as the false discovery rate (FDR) threshold to identify motifs showing motif importance significantly above the background level. Next, we split the region around the maximum cleavage site into 4 bins [(−120,−30), (−30,0), (0,30), (30,120)] and compared the 20 nt sum importance window score of each motif to the distribution of scores for other motifs in that same region. We retained those motifs with scores that exceeded the 99^th^ percentile in a region. This process identified 136 motifs that significantly contributed to PolyaClassifier predictions in *S. cerevisiae* and 182 motifs in *S. pombe*.

The significant motifs we identified in *S. cerevisiae* belong to three families: (1) the UA-rich family which contained motifs similar to UAUAUA or AUAUAU, (2) the A-rich family which are near-cognates of AAAAAA, and (3) the U-rich family which was represented by UUUUUU. We measured the motif similarity to each of the archetypical motifs using the Hamming distance, which is equal to the number of mismatches in the sequence (i.e., AUAUAA compared to AAAAAA has a Hamming distance of 2). Motifs were included in a family if they had a Hamming distance ≤2 nt. We then calculated the mean per-site and sum importance profiles for families of motifs, as shown in Figure 2AB.

*S. pombe* is distantly related to *S. cerevisiae* on an evolutionary time scale and is known to show divergent patterns of polyadenylation regulation (Liu et al. 2017). The contributing motifs we identified above belong to three unique families: (1) the A-rich family containing motifs with ≤3 mismatches compared to AAAAAA, (2) the U-rich family containing motifs with ≤2 mismatches compared to UUUUUU, and (3) the GUA-containing motif family. We calculated the mean per-site and sum importance profiles for these families of motifs, shown in Figure 6F and Figure S5A, as described above.

### Identifying the regulators of polyA site cleavage heterogeneity

Some genes in *S. cerevisiae* undergo extensive heterogeneous cleavage, while others show more precise cleavage (Ozsolak et al. 2010; Pelechano et al. 2013; Moqtaderi et al. 2013). We measured the degree of heterogeneity at a cleavage site using the entropy value. Using the PASS reads, we compiled the cleavage probability vector for the 50 nt window centered at the maximum cleavage site. Supposing the total read number from the polyA region is *n* and the read count in position *i* is *m*_*i*_, we calculated the fraction of reads *p*_*i*_ = *m*_*i*_/*n*. We then calculated the entropy value *E* using the cleavage vector distribution as follows:

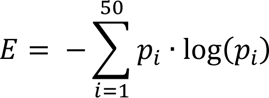

The entropy values were used to partition sites into three groups: low entropy sites that exhibited very defined cleavage sites (bottom 20% of entropy values, N = 3229), middle entropy sites (middle 60%, N = 9684), and high entropy sites that demonstrated high cleavage heterogeneity (top 20%, N = 3228).

To determine the motifs contributing to low versus high cleavage entropy, we split the sequences around each maximum cleavage site into 5 regions [(−90,−50), (−50,−30), (−30,−15), (−15,15), (15,30)]. For each region, we conducted a Chi-squared test to identify motifs enriched in the sequences from either high or low entropy group. We further confirmed these findings by comparing the motif counts across entropy groups. We also used the same approach to quantify the cleavage heterogeneity for polyA sites in *S. pombe* and humans.

### Developing the PolyaCleavage model

#### Input dataset

To determine whether primary polyA site sequences determine the cleavage heterogeneity, we sought to train a model that uses the 240-nt sequence as the input to predict the cleavage profile. We calculated the cleavage probabilities for the region +/−25 nt around the maximum cleavage sites using the PASS reads, as the ratio between the read number at each position and the total read count in the region. The 50-nt long vector represented the cleavage probability at each site and the sum is 1.

#### Model architecture and training

For model training, we sampled the top 20,000 expressed sites across genes and the 240 nt sequences centered at these cleavage sites were one-hot encoded as described above. The dataset was split into 80% training, 10% validation, and 10% testing sets. The model was trained using the Adam optimizer with Nesterov momentum to minimize the Kullback-Leibler divergence loss function:

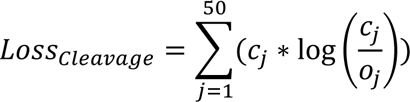

Where *c*_*j*_ and *o*_*j*_ are the predicted and observed cleavage probabilities, respectively, for each position j across the 50 nt window vector. The loss function was implemented out-of-box in TensorFlow2. The data flowed into the model using a batch size of 100 and a learning rate of 0.001. The training was stopped after 8 epochs when the minimum loss on the validation split was reached. Further details about the model architecture are in Table S2.

#### Performance evaluation

To evaluate the performance of the PolyaCleavage model, we compared the observed and predicted cleavage heterogeneity as well as mean cleavage position (MCP), which were calculated from the 3’READS cleavage probability vector and the predicted cleavage vector, respectively. The cleavage heterogeneity was quantified by the entropy score described above. The MCP is the dot product between the position indices and the cleavage probability vector, which must sum to 1. For position *i* from a vector of length 50 with a cleavage probability of *p*_*i*_, the MCP *m* was calculated as follows:

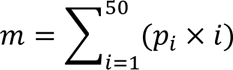

The resulting MCP measures the weighted mean and indicates the most likely cleavage position. The correlation between the observed and predicted entropy scores and MCP values quantifies the similarity between the 3’READS distribution and our model predictions surrounding a given polyA site.

### Examining genomic parameters regulating cleavage heterogeneity using the PolyaCleavage model

We found that both the nucleotide composition around the cleavage site and the presence of multiple UA-rich motifs upstream contribute to increased cleavage heterogeneity. To verify these findings, we sought to alter the cleavage entropy of low and high entropy sites by adding or removing these elements. We randomly added non-overlapping A-rich motifs into the (−15,+15) nt region around the cleavage site of low entropy sites. When introducing new motifs to the sequence, we added motifs that were identified through our motif enrichment analysis shown in Figure 3B and were most significant: UAAUAA, AUAAUA, AAUAAU, or AAACUA. We then measured the predicted changes to cleavage entropy using the PolyaCleavage model. Inversely, we randomly added non-overlapping U-rich motifs to the same region of high entropy sites. In this analysis, we introduced the top U-rich motifs that were significantly enriched in low entropy sites from our motif analysis: UUUUUU, UUCUUU, UCUUUU, UUUUUC, UUUCUU, UUUUCU, or CUUUUU. For each experiment, we highlighted representative examples by showing the modified sequence and predicted cleavage vector after each sequential motif addition.

We also modified the number of upstream efficiency elements to confirm the influence of these motifs on the cleavage heterogeneity. We sequentially disrupted the upstream efficiency elements of high entropy sites with five existing UA-rich motifs in the (−90,−30) region. We replaced the UA-rich element with randomly sampled nucleotides and repeated this disruption 100 times. We then measured the predicted change in cleavage entropy after modifying the sequence using PolyaCleavage.

### Developing the PolyaStrength model

#### Input dataset

As some nearby cleavage sites can result from the heterogeneous cleavage of the same polyA site, we clustered these closely located sites to quantify their expression levels. To this end, we grouped adjacent cleavage sites through an iterative clustering approach. We started by selecting the highest expressed site not yet assigned to a cluster. We searched 20 nt upstream and 20 nt downstream and added all cleavage sites within this region to the cluster. We repeated this process until all sites were clustered. We picked the 20 nt window because the A-rich motif (CF1A) located 20-nt upstream is important for the downstream cleavage. We considered that the sites located within 20 nt of a maximum cleavage site are likely due to heterogeneous cleavage. If the boundaries of the two clusters were within 10 nt of each other, we adjusted the cluster boundaries to evenly split the area between the maximum cleavage site of each cluster. For example, if the maximum sites of two neighboring clusters were 8 nt apart, the edges of the cluster would be adjusted so that each cluster covered 4 nt of that region. We filtered out clusters supported by fewer than 10 3’READS or with expression < 2% of the maximum number in each gene. This process yielded 29,430 polyA site clusters in *S. cerevisiae*.

We selected clustered polyA sites within the 3’UTR or extended downstream region of genes undergoing APA and therefore contained at least 2 expressed polyA sites supported by 3’READS (N = 18,434 sites). These sites formed the dataset for the PolyaStrength model. For genes containing multiple polyA sites, we calculated the relative usage of each site as measured by the PASS read number assigned to the site to the sum of reads supporting the top two sites in a gene. Supposing the supporting read number for site *i* is *n*_*i*_, its relative usage *u*_*i*_ and the logit-transformed value *o*_*i*_ were calculated as follows:

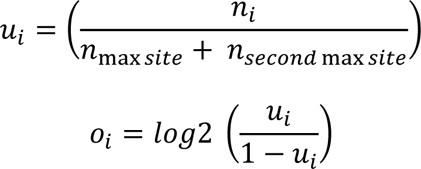

The maximum cleavage site within each clustered polyA site served as the representative site for subsequent model training and analysis.

#### Model architecture and training

We used the 240 nt sequence surrounding the representative cleavage site of each cluster as the input for model training. The sequences were one-hot encoded as described above. The PolyaStrength model architecture is similar to the PolyaClassifier model, but the parameters vary slightly and are described in detail in Table S2. The output layer uses a linear activation to predict the logit-transformed usage levels. The dataset was split into 80% training, 10% validation, and 10% testing data splits. The model was trained using the Adam optimizer with Nesterov momentum to minimize the root-mean-squared error:

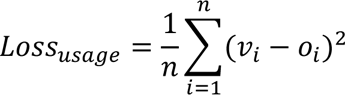

Where *o*_*i*_ and *v*_*i*_ are the observed and predicted logit-transformed usage levels for the polyA site *i*. The loss function was implemented out-of-box in TensorFlow2. The data flowed into the model using a batch size of 32 and an initial learning rate of 0.001. After the 2^nd^ training epoch, we decreased the learning rate by a factor of *e*^−0.1^ each subsequent epoch. The training was stopped after 7 epochs when the minimum loss on the validation split was reached.

#### Performance evaluation

To evaluate the utility of the PolyaStrength model, we created pairs of polyA sites from the holdout test set located in the same 3’UTR. We required that the paired sites have a usage difference >8-fold, be ≥25 nt apart, and were not included in the model training. 495 pairs of sites were created for S. cerevisiae. We then calculated the AUROC and AUPRC using the difference in predicted PolyaStrength scores to classify the stronger vs. weaker of a pair.

### Identifying motifs contributing to polyA site strength

We used the similar hexamer disruption and motif analysis approaches used above for the PolyaClassifer model to identify motifs driving the polyA strength. We used 7,378 clustered polyA sites with ≥100 reads and ≥5% usage for this analysis. 184 motifs were found to show significantly higher sun importance scores than expected (Table S5). These motifs were similar to what we found driving polyA site formation by analyzing the PolyaClassifer model.

### Stuyding the APA regulation under diauxic stress

We selected the top two expressed polyA site clusters in each gene by pooling the 3’READS measured under rich media and diauxic stress culture conditions. We then defined the polyA site clusters using a similar approach we used above for calculating the polyA site strengths. We selected genes showing significant APA under the stress condition comparing proximal vs. distal sites using the cutoff Benjamini-Hochberg correction *P*-value <0.05 (Chi-squared test). For these genes, we calculated the proximal polyA site usage levels as the ratio between the PASS read number supporting proximal sites vs. the sum reads of both proximal and distal. We next grouped polyA sites based on their differential usage levels between the diauxic stress vs. rich media conditions. We characterized these sites using the PolyaStrength scores, the difference in PolyaStrength between the proximal and paired distal sites, and the distance between the two sites.

#### Examing the context of GUA-elements in *S. pombe*

To quantify the importance of the nucleotides surrounding GUA elements in *S. pombe* and determine their most significant context, we padded “GUA” with all combinations of 3 nucleotides on each end to create 4096 possible 9-mers with GUA in the center. We then applied our systematic disruption approach to quantify the importance of each GUA-containing 9-mer where the GUA was found (15,60) nt downstream of the maximum cleavage site. For each position around the central GUA element, we grouped the motifs containing each nucleotide and calculated the mean importance score. These scores were combined and normalized to sum to 1 to create a position-probability matrix (PPM). We calculated the log-likelihood of the PPM assuming that all nucleotides are equally important and plotted the position-weight matrix for the nucleotides with positive values.

### Comparing polyA site conservation across species

To examine the sequence conservation surrounding polyA sites across yeast and humans, we used the PhastCons conservation tracks (Siepel et al. 2005). For humans, we used the 100-way alignment from UCSC. For *S. cerevisiae*, we used the 7-way alignment compared to other *Saccharomyces* species. For *S. pombe*, we used the 4-way alignment from the Fungal Genome Initiative comparing *S. pombe* to other *Schizosaccharomyces* family members (Rhind et al. 2011). We quantified the mean conservation score within coding regions and used this as a normalization factor. We performed the analyses for homologous genes across the three species defined by the PomBase database (Harris et al. 2022). We selected the top expressed polyA site from each homologous protein-coding gene (N = 6546 for *S. cerevisiae*, 6373 for *S. pombe*, and 8170 for *H. sapiens*). We calculated the averaged conservation scores and the 95% confidence interval values at each position normalized to the mean coding region conservation. We also performed similar analyses for polyA sites from non-homologous genes, and sites grouped based on their relative genomic locations.

## ACKNOWLEDGMENTS

This work was supported by grants to Z.J.: the National Institutes of Health (R35GM138192, and R01HL161389), and the Lynn Sage Scholar fund. E.S. was supported by the Predoctoral Training Program in Biomedical Data Driven Discovery (T32LM012203). We thank the members of the Ji lab for helpful discussions.

## DATA AVAILABILITY

The sequencing datasets analyzed in this study are available in the Gene Expression Omnibus (GEO) repository with the accession number GSE67163, GSE95139, and GSE151196.

## CODE AVAILABILITY

The source codes were deposited in GitHub: https://github.com/zhejilab/PolyaModelsYeast.

## SUPPLEMENTARY FIGURES

**Supplemental Figure 1.**
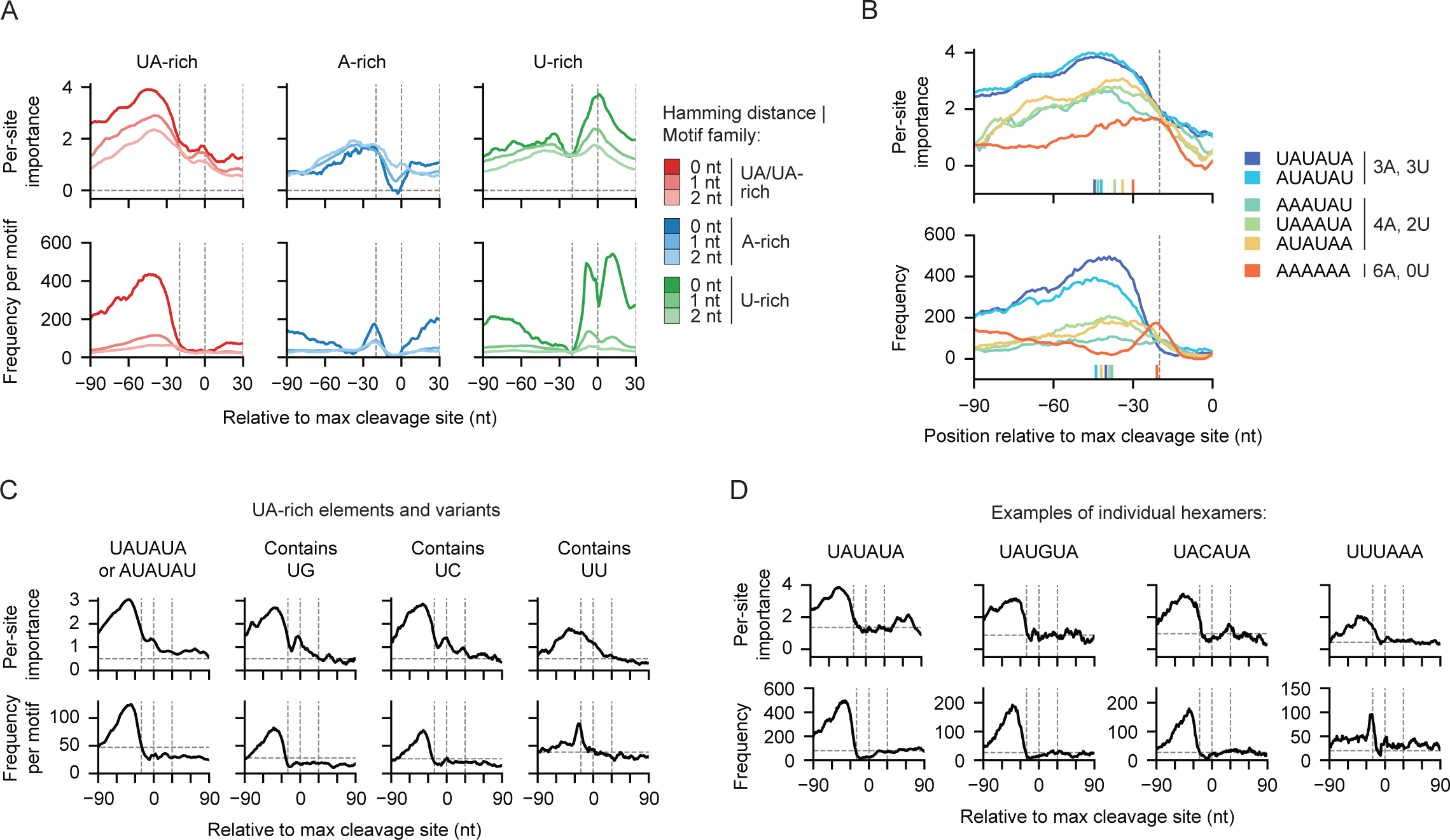
Examine motif importance to polyA site identification by motif family. (A) The per-site importance (top) and frequency (bottom) profiles for the *S. cerevisiae* motif families. In each family, the importance profile is grouped by the Hamming distance to the representative motif (0, 1, or 2 nt). (B) Examples of per-site importance (top) and frequency (bottom) for individual motifs grouped by the number of A and Us. The dashes along the x-axis indicate the location of the maximum for each motif. (C) The per-site importance (top) and frequency (bottom) profiles for the UA-rich motifs. Motifs are grouped by whether they contain nucleotide variants. (D) The per-site importance (top) and frequency (bottom) of example UA-rich motifs.

**Supplemental Figure 2.**
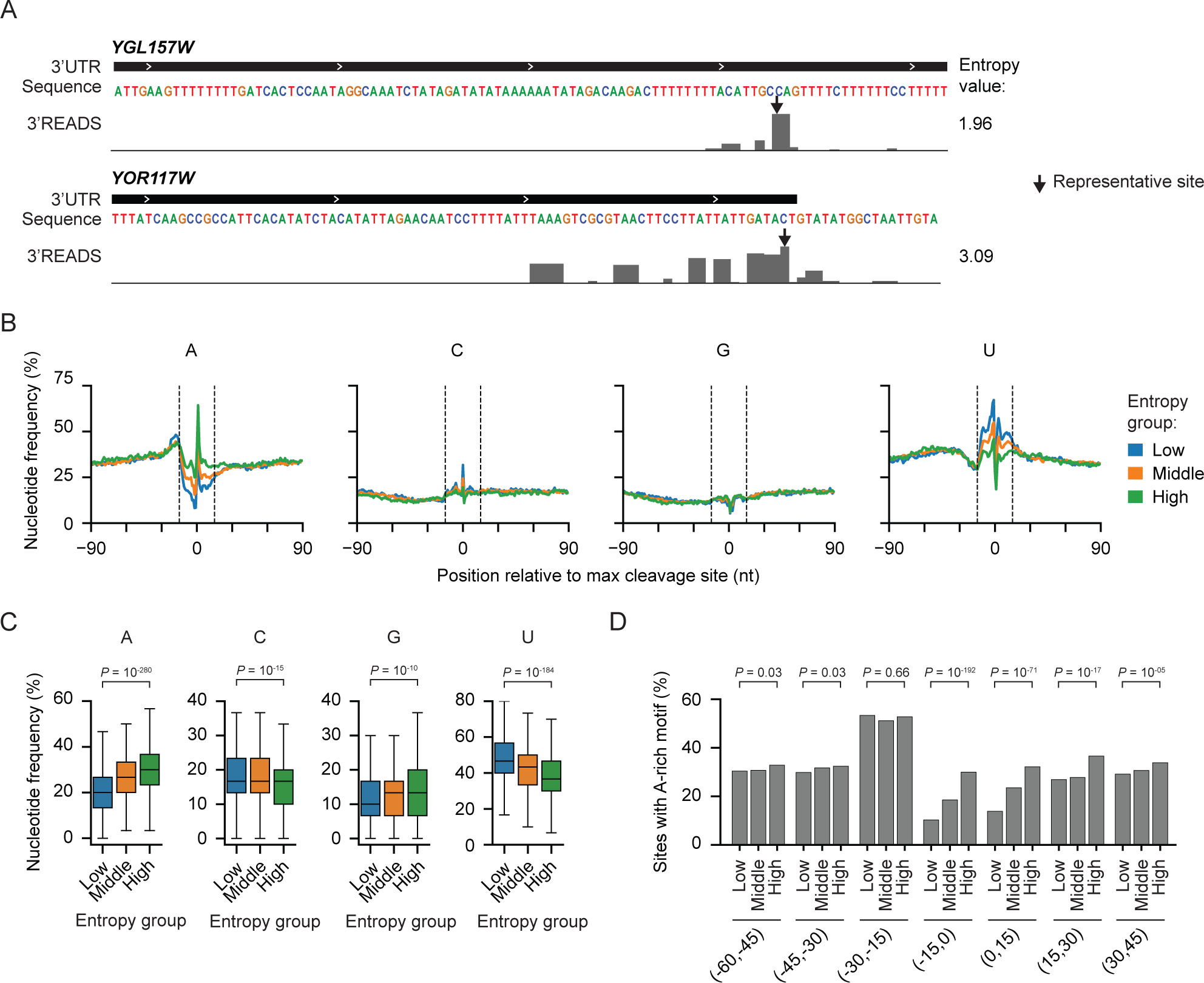
Motif analyses of polyA sites from entropy groups. (A) Examples of the PASS read distribution surrounding a low entropy site (top) and high entropy site (bottom). The observed entropy values are shown. (B) The nucleotide frequency surrounding the maximum cleavage site grouped by cleavage entropy. The vertical dashed lines mark −15 nt and +15 nt. (C) The quantification of nucleotide density in the (−15,+15) nt region grouped by cleavage entropy. The *P*-values from the Wilcoxon rank-sum test comparing low vs. high entropy sites are shown. (D) The fraction of sites in each cleavage entropy group that contain an A-rich motif in the noted regions. The *P*-value from the two proportions hypothesis test is shown.

**Supplemental Figure 3.**
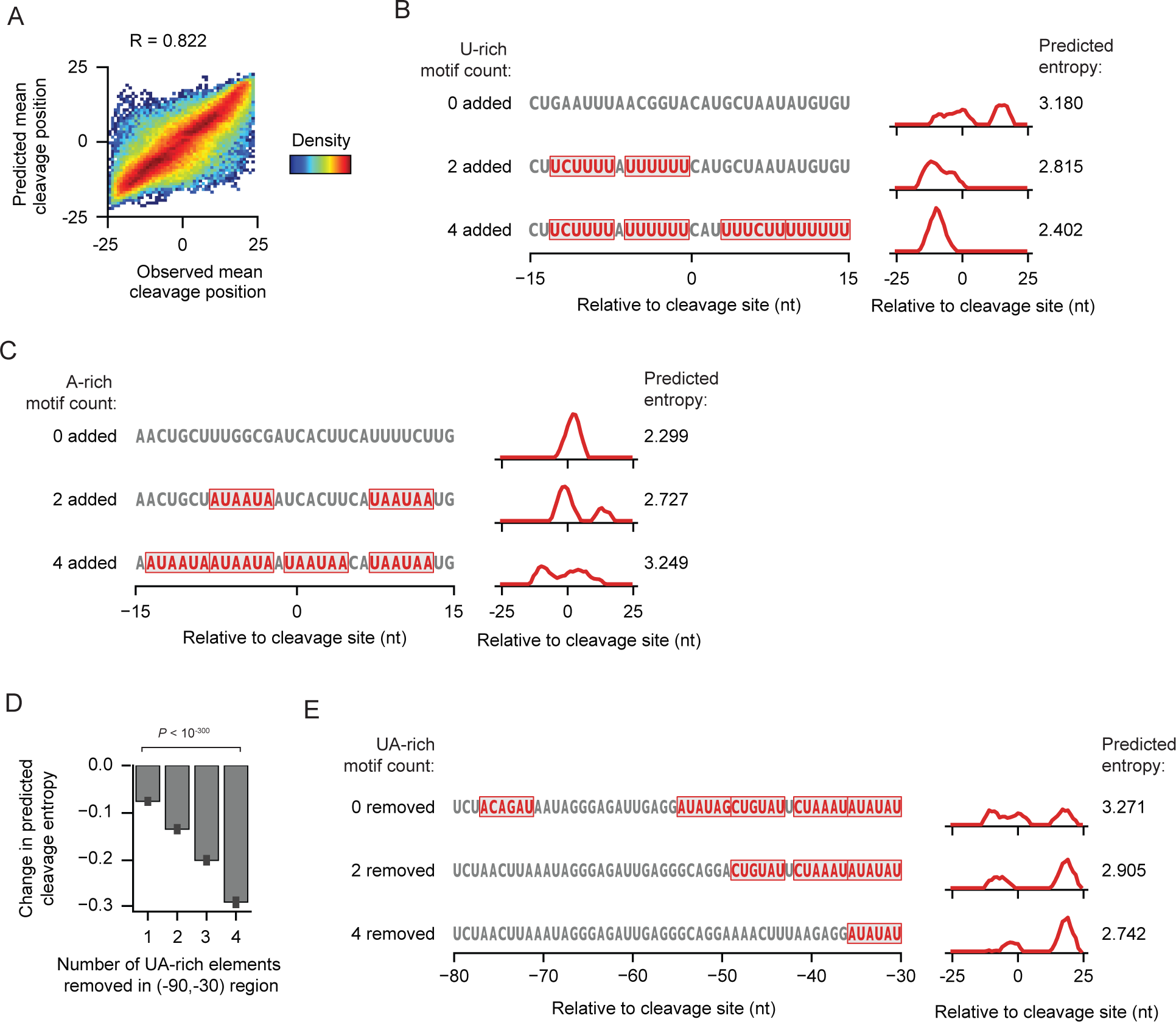
Examples of cleavage site entropy conversions. (A) A 2D heatmap showing the correlation between the observed and predicted mean cleavage position in the holdout testing set (N = 102001). The Pearson correlation coefficient is shown. (B) An example showing the effect of adding U-rich motifs in the (−15,+15) nt region around the max cleavage site in the *YJR118C* gene. (C) An example showing the effect of adding A-rich motifs in the (−15,+15) nt region around the max cleavage site in the *YLR330W* gene. (D) The change in predicted cleavage entropy as upstream UA-rich elements are sequentially disrupted in the (−90,−30) region. High entropy sites with 5 upstream UA/UA-rich motifs in this region were included. The data is shown as the mean and the 95% confidence interval (error bar). The *P*-value from the Wilcoxon rank sum test comparing sequences with 1 motif removed vs. 4 motifs removed is shown. (E) An example showing the effect of disrupting upstream UA-rich motifs in the YER120W gene.

**Supplemental Figure 4.**
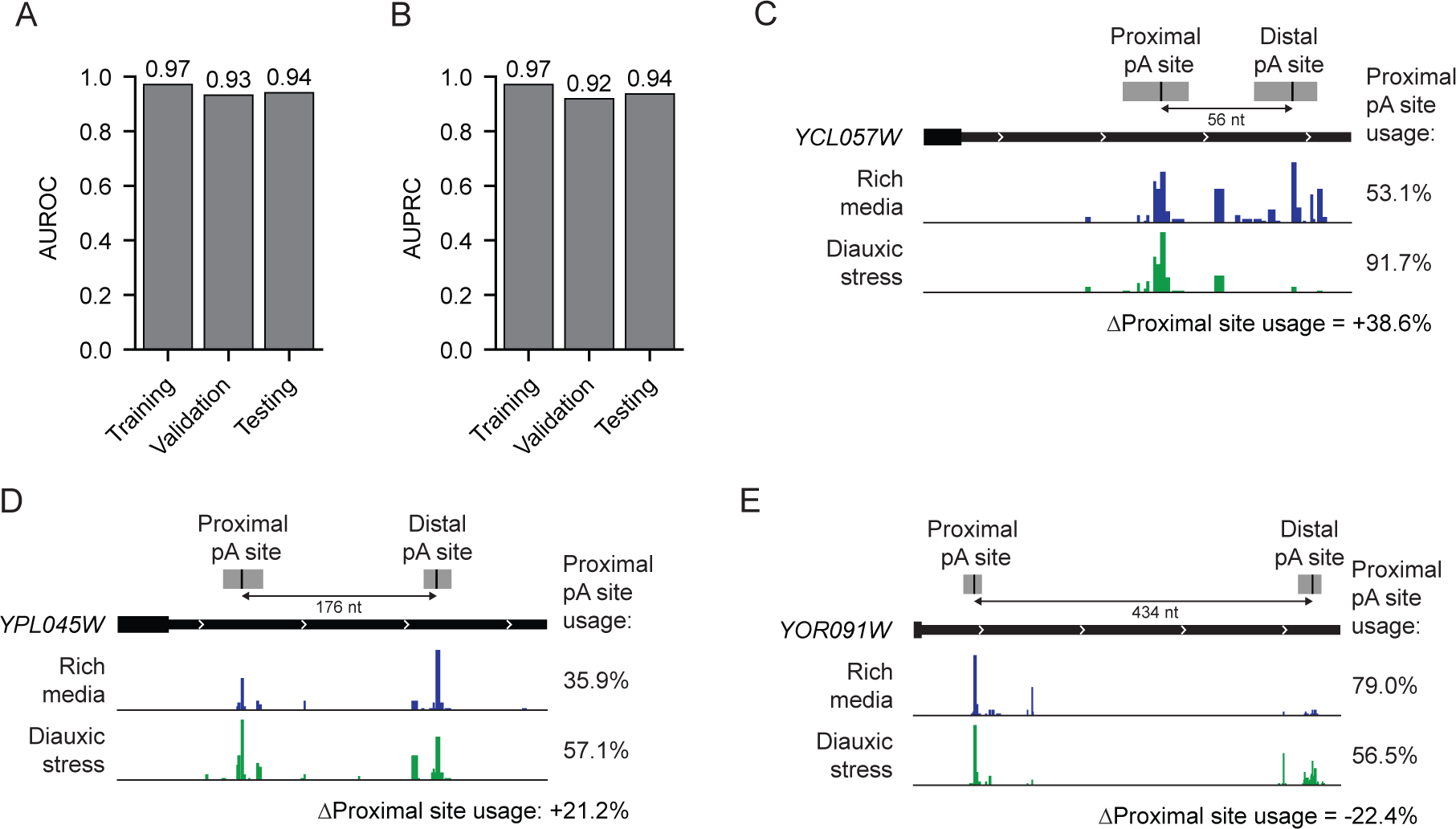
The evaluation of PolyaStrength model performance and examples of APA regulation under the diauxic stress. (A) The AUROC values for the training, validation, and testing datasets for the *S. cerevisiae* PolyaStrength model. (B) Similar to (A), except showing the AUPRC values. (C) An example gene *YCL057W* that shows increased proximal site usage under diauxic stress. The proximal and distal polyA site clusters are shown, with the distance between them noted. The percentage usage levels of the proximal site are shown. (D) Similar to (C), showing the *YPL045W* gene. (E) Similar to (C), the *YOR091W* gene shows decreased proximal site usage under diauxic stress.

**Supplemental Figure 5.**
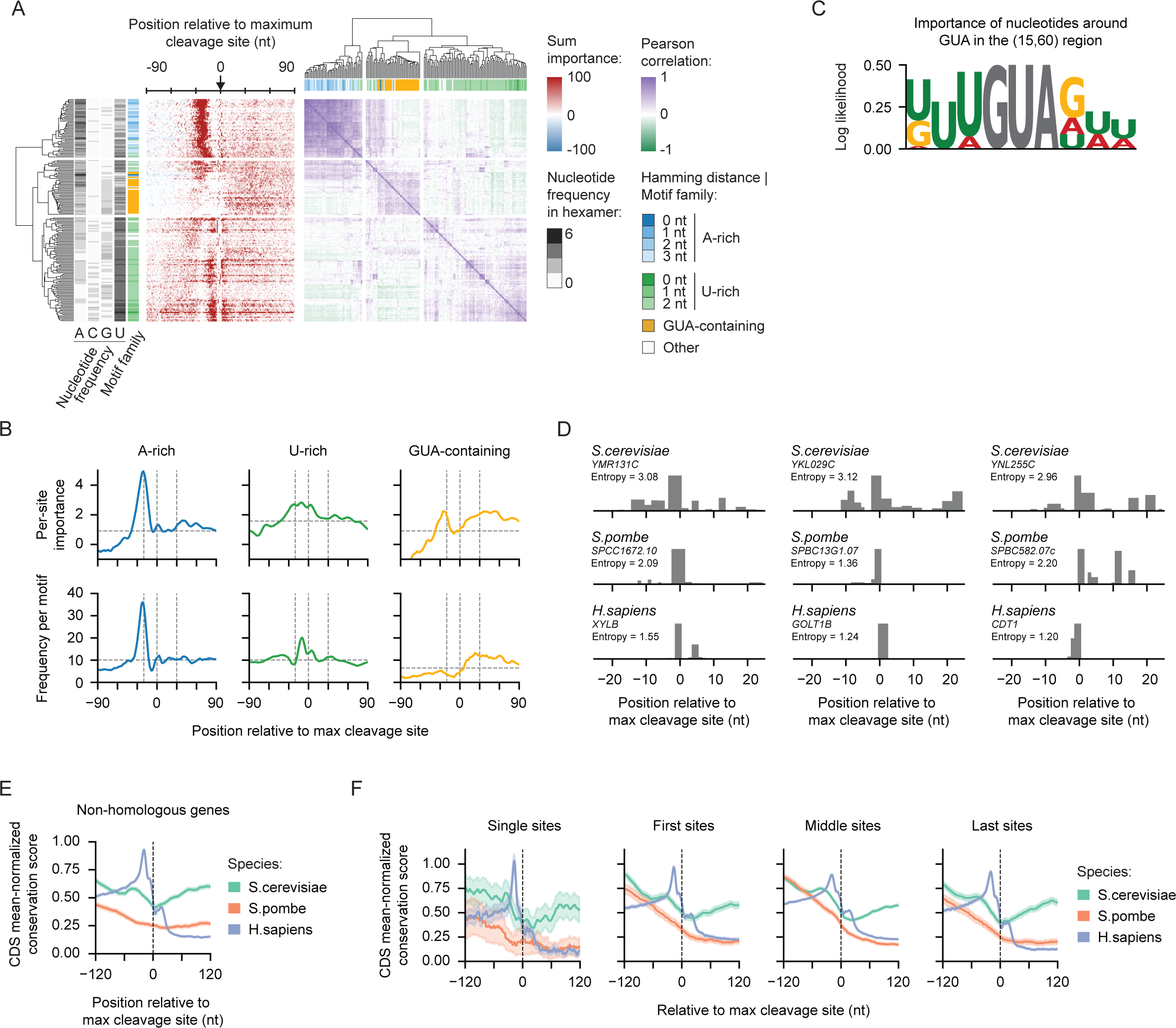
Anlyases of *S. pombe* polyA site motifs and site conservation. (A) Heatmaps showing the sum classification importance (left) and Pearson correlation between classification importance profiles (right) for *cis*-regulatory elements significantly contributing to polyA site definition in *S. pombe* (N = 182). Each row represents a hexamer. The A-rich and U-rich motif families are defined by the Hamming distance between the motif of interest and the archetypical motifs AAAAAA or UUUUUU, respectively. A third family for motifs containing GUA is also shown. (B) The per-site importance (top) and frequency (bottom) profiles for A-rich, U-rich, and GUA-containing motifs that significantly contribute to polyA site definition in *S. pombe*. (C) A position-weight matrix showing the log likelihood of the importance of nucleotides surrounding GUA 3-mers in the (15,60) region. See Methods for detailed calculation. (D) Additional examples of the PASS read distribution surrounding the top polyA site in homologous genes across *S. cerevisiae, S. pombe,* and *H. sapiens*. The observed cleavage entropy values are shown. (E) A meta plot showing the conservation score for the +/−120 nt surrounding the top expressed polyA sites in all non-homologous genes across the three species. The conservation score was normalized to the mean score in coding regions. The data is shown as the mean with the shaded region indicating the 95% confidence interval. (F) Similar to (E), except for well-expressed polyA sites grouped by their relative position in the gene.

## SUPPLEMENTARY TABLES

**Supplemental Table 1. The 3’READS datasets analyzed in this study.**

**Supplemental Table 2. The detailed architectures of deep learning models we developed in this study.**

**Supplemental Table 3. The motifs showing significant importance scores from the PolyaClassifer model in *S. cerevisiae*.**

**Supplemental Table 4. The polyA sites identified in *S. cerevisiae*.**

**Supplemental Table 5. The motifs showing significant importance scores from the PolyaStrength model in *S. cerevisiae*.**

**Supplemental Table 6. The polyA sites identified *in S. Pombe*.**

**Supplemental Table 7. The motifs showing significant importance scores from the PolyaClassifer model in *S. Pombe*.**

